# Reduced recruitment of inhibitory control regions in very young children with ADHD during a modified Kiddie Continuous Performance Task: a fMRI study

**DOI:** 10.1101/2024.01.17.576033

**Authors:** Mohammadreza Bayat, Melissa Hernandez, Madeline Curzon, Dea Garic, Paulo Graziano, Anthony Steven Dick

## Abstract

Attention-Deficit/Hyperactivity Disorder (ADHD) symptom profiles are known to undergo changes throughout development, rendering the neurobiological assessment of ADHD challenging across different developmental stages. Particularly in young children (ages 4 to 7 years), measuring inhibitory control network activity in the brain has been a formidable task due to the lack of child-friendly functional Magnetic Resonance Imaging (fMRI) paradigms. This study aims to address these difficulties by focusing on measuring inhibitory control in very young children within the MRI environment. A total of 56 children diagnosed with ADHD and 78 typically developing (TD) 4-7-year-old children were examined using a modified version of the Kiddie-Continuous Performance Test (K-CPT) during BOLD fMRI to assess inhibitory control. We concurrently evaluated their performance on the established and standardized K-CPT outside the MRI scanner. Our findings suggest that the modified K-CPT effectively elicited robust and expected brain activity related to inhibitory control in both groups. Comparisons between the two groups revealed subtle differences in brain activity, primarily observed in regions associated with inhibitory control, such as the inferior frontal gyrus, anterior insula, dorsal striatum, medial pre-supplementary motor area (pre-SMA), and cingulate cortex. Notably, increased activity in the right anterior insula was associated with improved response time (RT) and reduced RT variability on the K-CPT administered outside the MRI environment, although this did not survive statistical correction for multiple comparisons. In conclusion, our study successfully overcame the challenges of measuring inhibitory control in very young children within the MRI environment by utilizing a modified K-CPT during BOLD fMRI. These findings shed light on the neurobiological correlates of inhibitory control in ADHD and TD children, provide valuable insights for understanding ADHD across development, and potentially inform ADHD diagnosis and intervention strategies. The research also highlights remaining challenges with task fMRI in very young clinical samples.

## 1| INTRODUCTION

Externalizing behavior problems, including symptoms of attention-deficit/hyperactivity disorder (ADHD) such as inattention, hyperactivity, and impulsivity, are the most common reasons for early childhood mental health referrals [1, 2, 3, 4]. These symptoms are highly prevalent during the preschool and early elementary period (ages 4-7 years) and can be challenging to assess [5]. Moreover, the symptom profile of young children with ADHD can significantly change over their developmental course [6], indicating the need for measures that can be appropriately employed from preschool into later adolescence and adulthood. Despite the existence of some lab-based behavioral measures, there is a dearth of neuroimaging studies focusing on young children with ADHD [7, 8, 9, 10, 11, 12, 13, 14], and even fewer that utilize well-developed magnetic resonance imaging (MRI) task paradigms [11]. This scarcity can be attributed to the challenges of designing tasks that assess the behaviors of interest in an MRI environment, particularly with very young children who must remain still and attentive for an adequate duration. The lack of such tasks contributes to inconsistent measurements of ADHD-related behaviors and hampers researchers’ understanding of the disorder’s development.

While addressing the need for consistent measurement remains a challenge, it is essential for several reasons. First, defining ADHD poses a persistent challenge due to the diversity of symptom profiles among affected children, who experience impairments across various domains. For instance, inhibitory control is consistently impaired in children with ADHD [15], but not all individuals with ADHD exhibit this deficit, as up to 25 percent of diagnosed children do not show such impairments [16]. Notably, these studies are primarily conducted in older children, and the disorder’s profile changes significantly as children progress through childhood. Consequently, it is crucial to track the nature of ADHD heterogeneity over development, which is complicated by the lack of reliable measures in young children. A second imperative for consistent measurement lies in improving the accuracy of prevalence estimates. Variability in research methodologies, broad age ranges assessed, and changes in diagnostic criteria over time can lead to divergent estimates of ADHD prevalence [6]. Lastly, achieving better measurement of the neurobiological profile of ADHD in early childhood is vital to defining the disorder at multiple levels of analysis. Although there is consensus that certain areas involved in inhibitory control, such as the frontal, parietal, basal ganglia, and cerebellum, are affected in ADHD, the neurobiological definition of the disorder remains insufficiently established. These brain regions also play roles in cognitive and affective processes often impaired in children with ADHD, such as executive function and emotion regulation [14]. Addressing definitional issues at multiple levels of analysis is crucial for understanding ADHD’s etiology, especially in very young children. However, current methods do not adequately support this endeavor, as there are limited well-established functional imaging tasks for young children. Researchers typically rely on electroencephalography and functional near-infrared spectroscopy (fNIRS) [17, 18, 19, 20, 21, 22] to investigate these questions, but these methods lack the spatial resolution of fMRI, preventing them from addressing the specific questions that functional MRI could answer.

## 2| DEVELOPING INHIBITORY CONTROL IN ADHD

In this study, our primary objective is to address the challenges associated with measuring inhibitory control in the MRI environment, specifically focusing on very young children aged between 4 and 7 years. Inhibitory control, as defined by Aron and colleagues [23], involves the ability to suppress inappropriate responses, stimulus-response mappings, or task-sets when the context changes, as well as the suppression of interfering memories during retrieval. This is typically measured in adults using tasks that require the suppression of a prepotent response that is established by the rules of the task. For example, in stop-signal paradigms [24] a participant must respond to a target (usually via a button press), but withhold responding when a stop signal appears. Withholding the response requires inhibitory control. A similar paradigm used is the Go/NoGo paradigm [25, 26, 27, 28, 29, 30], where participants must press a button when presented with “Go” stimuli, but refrain from responding when encountering a “NoGo” stimulus.

Variations on this paradigm are often called “Continuous Performance Tasks” (CPT) because of the need to maintain attention over the course of the stimulus presentation [31]. In fact, children with ADHD often perform more poorly on CPT tasks than typically developing (TD) children. Notably, the Kiddie Continuous Performance Task (K-CPT) [32] is a well-established measure of attention maintenance and inhibitory control, widely utilized in both laboratory and clinical settings with very young children. Unlike some CPT paradigms that involve responding to rare signals, the K-CPT is more akin to inhibitory control tasks. In this task, children first establish a prepotent response (e.g., pressing a button upon seeing a picture), and then must inhibit this response when presented with a “NoGo” signal (e.g., a picture of a soccer ball).

Previous studies have demonstrated that the K-CPT administered in the laboratory or clinic is highly sensitive in distinguishing children with ADHD from TD children [33, 34, 35, 36]. For instance, Breaux and colleagues [37] found that among various neuropsychological measures, the K-CPT was the most effective in predicting whether young children with high externalizing behaviors at age 3 would later be diagnosed with ADHD at age 6. In addition to behavioral assessments, researchers have explored the neural underpinnings of inhibitory control during the K-CPT using electroencephalography (EEG) [38, 39].

Despite the effectiveness of the K-CPT in distinguishing ADHD from TD children, no studies have investigated the specific brain regions differentially engaged in young children with ADHD compared to TD children, as fMRI paradigms for the K-CPT in these groups have not been established. In fact, few studies have attempted to assess inhibitory control in children with ADHD, and at any rate these have used different paradigms and focused on children aged above 7 years [40, 41, 42, 43, 44, 45, 46, 47, 48]. In a study of 6-14-year-olds using a CPT paradigm with fNIRS, Inoue and colleagues [22] found reduced activation in frontal regions in the ADHD group during conditions with high inhibitory control demand. These results suggest reduced recruitment for the ADHD group, but the fNIRS paradigm does not have the spatial resolution to indicate what parts of this broad network are recruited to a reduced extent, or indeed what specific regions comprise that network in very young children.

However, despite the lack of specific fMRI paradigms for the K-CPT in young children, there are expectations about the neural networks involved in inhibitory control during CPT tasks, largely derived from studies conducted in adults [49, 50, 51, 52]. Models of the neurobiology of inhibitory control have been proposed, such as the computational model of inhibitory control by Wiecki and Frank [30], and the anatomical model of stopping by Aron and colleagues [53]. These models suggest that inhibitory control in CPT paradigms relies on a specific network of cortico-basal ganglia regions, including the right hemisphere’s inferior frontal gyrus (IFG) and neighboring anterior insula, the presupplementary motor area (pre-SMA), the dorsal striatum, the subthalamic nucleus (STN), and other basal ganglia regions.

Within this network, specific brain regions contribute in different ways. The subthalamic nucleus, emphasized by Aron and colleagues [53], plays a crucial role in outright action stopping and collaborates with the IFG/anterior insula and pre-SMA, which may trigger STN activity via a hyperdirect pathway [53]. It is anticipated that these regions may function differently in children with ADHD compared to TD children, and the K-CPT administered during fMRI may reveal functional activation differences in these particular brain areas. Figure 1 highlights these regions, which will be a focal point in the present study.

**FIGURE 1.**
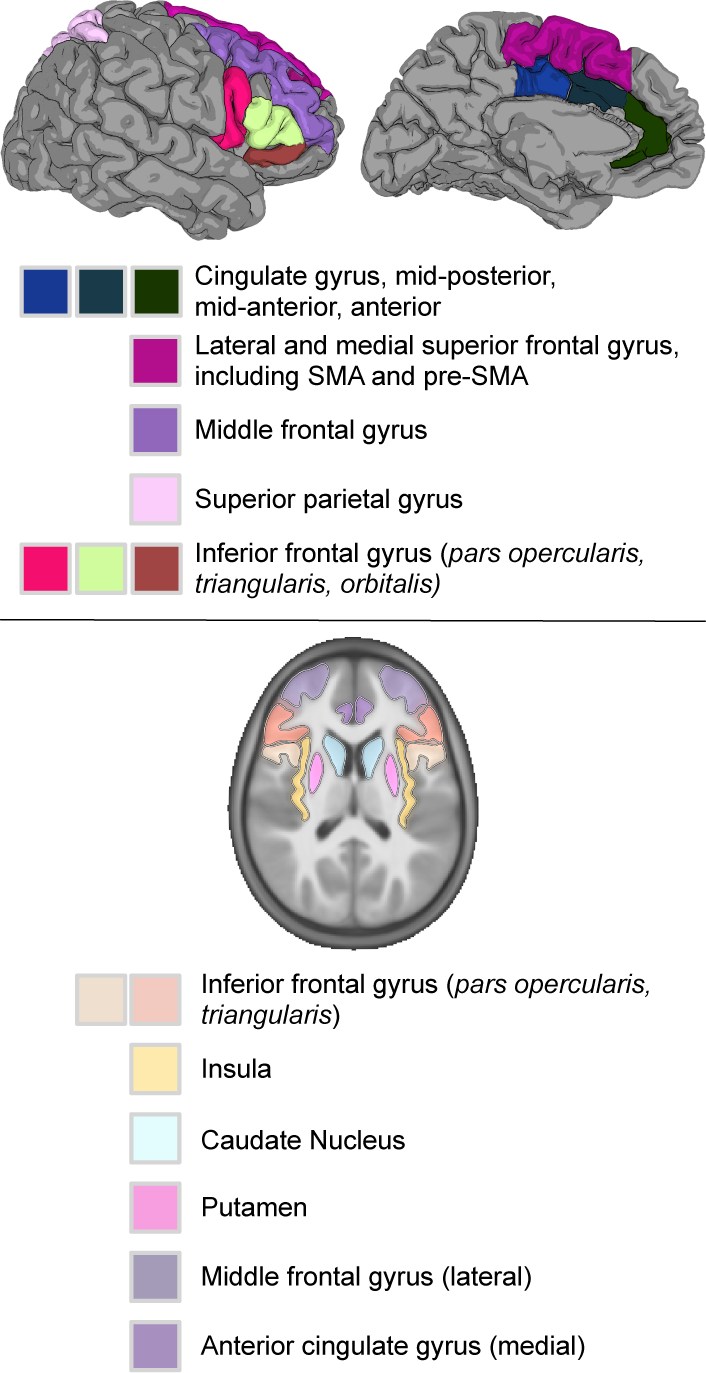
*Regions comprising the inhibitory control network.* Top: Cortical surface representation shows regions of lateral inferior frontal gyrus, superior parietal gyrus, and lateral middle and superior frontal gyri (i.e., dorsolateral prefron cortex), and medial superior frontal (including SMA and pre-SMA), and middle and anterior cingulate gyrus regions. Bottom: ABCD atlas representation in axial view also shows subcortical regions including caudate and putamen of the dorsal striatum, and the anterior insula neighboring inferior frontal gyrus (not visible from lateral view).

In summary, the present study aims to achieve three main objectives. First, we seek to evaluate the suitability of the K-CPT paradigm in identifying the potential network involved in inhibitory control among very young children (ages 4-7 years) with and without ADHD. Secondl, we aim to explore the differential recruitment of this network in young children with ADHD compared to typically developing (TD) children. Lastl, we aim to establish connections between performance on the K-CPT and the response profile in the inhibitory control/executive function network, and analyze how these profiles vary across the two groups.

Our predictions are as follows: First, both groups are expected to engage a broad cortical and subcortical network involved in inhibitory control, as illustrated in Figure 1, to perform well on the K-CPT. Secondl, children with ADHD are anticipated to exhibit reduced activation in the identified inhibitory control regions, indicating a main effect of group and indicating reduced recruitment of the network. Lastl, task performance will correlate with increased activation in the identified inhibitory control regions; however, this association is expected to differ between the two groups, suggesting a group-by-brain activity interaction when predicting performance.

## 3| MATERIALS AND METHOD

### 3.1| Participants

The proposed final sample included 56 children diagnosed with ADHD and 78 typically developing (TD) children. An additional 193 children were recruited as part of a broader study on ADHD, but 31 of them failed to complete either the T1-weighted scan or the experimental task EPI scan, 129 exhibited excessive movement during the T1-weighted MRI or task EPI, exceeding pre-determined thresholds (see Movement below), and 33 were identified as left-handed according to the Edinburgh Handedness inventory. Therefore, these 194 children were excluded from the current analysis.

To ensure consistent activation profiles across children, we restricted the analysis to right-handed participants as the task required a button press using the right hand. Prior to participation, each child and their parent provided written informed consent, following the guidelines of the Institutional Review Board for the Division of Social and Behavioral Sciences of Florida International University, which approved the study. Verbal assent was obtained from the children.

### 3.2| Recruitment and Exclusion Criteria

Participants and their caregivers were recruited through brochures, open houses, and parent workshops at local schools and mental health agencies. For the ADHD sample, parents were invited to participate in an assessment to determine study eligibility if they endorsed clinically significant levels of ADHD symptoms (six or more symptoms of either Inattention or Hyperactivity/Impulsivity according to DSM-5 or a previous diagnosis of ADHD), indicated that the child is currently displaying clinically significant academic, behavioral, or social impairments (score of 3 or higher on a seven-point impairment rating scale), and the child was not taking any psychotropic medication.

For the typically developing sample, parents and children were invited to participate in an assessment to determine study eligibility if they endorsed fewer than 4 ADHD symptoms (across either Inattention or Hyperactivity/Impulsivity according to DSM-5), fewer than 4 Oppositional Defiant Disorder (ODD) symptoms, and indicated no clinically significant impairment (score below 3 on the impairment rating scale).

Participants were also required to have been enrolled in school during the previous year, have an estimated IQ of 70 or higher, have no confirmed history of an Autism Spectrum Disorder, and for the ADHD group only, be able to attend an 8-week summer treatment program prior to the start of the next school year. Due to the young age of the sample, only disruptive behavior disorders were extensively examined for diagnostic purposes.

During intake, ADHD diagnosis and comorbid disruptive behavior disorders were assessed through a combination of parent structured interview [54] and parent and teacher ratings of symptoms and impairment [55] as is recommended practice [56]. Specifically, the DBD rating scales and diagnostic interview were combined using an “or rule,” which identifies the presence of a symptom if endorsed by either informant while clinically significant problems at home and school were defined by at least a “3” on a “0 to 6” impairment rating scale [57, 58]. A dual Ph.D. level clinician review was used to determine the diagnosis. The demographics of the final sample are shown in Table 1.

**TABLE 1.** Table of Demographics by Diagnostic Group.

|  | TD | ADHD | p-value |
| --- | --- | --- | --- |
| Sex assigned at birth (female) | 39.7% | 33.9% | 0.45 |
| Age | 5.45 yrs | 5.71 yrs | 0.07 |
| Parental education | 4.92 | 4.53 | 0.11 |
| T1 quality | 3.71 | 3.71 | 0.98 |
| Movement | 12.08 | 14.71 | 0.10 |

*Note.* Parental education and T1 quality were measured on 6-point ordinal scales. Movement was measured as the number of censored TRs.

### 3.3| Image Acquisition

All imaging was performed using a research-dedicated 3 Tesla Siemens MAGNETOM Prisma MRI scanner (V11C) with a 32-channel coil located on the university campus. Children first completed a preparatory phase using a realistic mock scanner in the room across the hall from the magnet. They were trained to stay still, and were also acclimated to the enclosed space of the magnet, to the back projection visual presentation system, and to the scanner noises (in this case, presented with headphones). Children were also trained on the fMRI tasks for the scanner, including the K-CPT (described below). Two other short fMRI tasks, and a diffusion-weighted imaging scan, were also acquired but are not analyzed here. When they were properly trained and acclimated, they were moved to the magnet.

Structural MRI scans were acquired using a 3D T1-weighted inversion-prepared RF-spoiled gradient echo sequence with 1 x 1 x 1 mm resolution, lasting 7 minutes and 14 seconds with prospective motion correction [59], according to the Adolescent BrainCognitive Development (ABCD) protocol [60]. The T2*-weighted echo-planar images optimized for blood oxygenation level-dependent (BOLD) effects were obtained with 2.5 mm isotropic resolution and 56 axial slices, using a repetition time (TR) of 1000 ms and an echo time (TE) of 30 ms, with an acceleration factor of 4. The total scan time was under 30 minutes.

### 3.4| Experimental paradigm

The experimental paradigm was a modified continuous performance test based on the Conners Kiddie Continuous Performance Test (K-CPT; [32]). Stimuli were projected onto a screen behind the MRI magnet, visible to the participants via a mirror attached to the imaging headcoil. The stimuli, which were taken from the K-CPT stimulus set, included pictures of various objects such as a bicycle, car, fish, hand, horse, house, sailboat, scissors, telephone, train, and soccer ball. Each picture was displayed for either 3000ms or 1500ms, with a 500ms interstimulus interval (ISI). During the ISI and between epochs, a red fixation cross was shown on a black screen.

The fMRI task followed a block design with four epochs, utilizing EPrime software (version 2.0.10.356 or later). The presentation was set to start with a trigger pulse from the MRI scanner. The task started with 30 seconds of fixation (with a 10-second pad for later censoring), followed by 36 seconds of continuous stimulus presentation. In each epoch, the soccer ball picture was randomly interspersed four times, resulting in a total of 16 soccer ball presentations across all epochs. Each epoch was separated by 20 seconds of fixation, and the block design concluded with 20 seconds of fixation, making a total of 254 TRs (254 seconds). The children were instructed to press a button as quickly and accurately as possible in response to any picture except the soccer ball, for which they were instructed to withhold their response. Prior to the actual MRI scanning, the children practiced the task in the mock scanner to ensure they understood the instructions. Response compliance was actively monitored during scanning, and omission errors (i.e., failing to hit the key when a target stimulus is presented), commission errors (i.e., hitting a key when the non-target soccer ball is presented), RT, and standard deviation of RT were recorded via the button box.

We also administered the standardized K-CPT 2nd Edition [32] outside the MRI magnet, which served as our primary behavioral measure of task performance because it is a standardized and validated measure. This version is administered on a computer and scored with accompanying software. On the K-CPT, children are presented with a series of pictures, the same stimuli that were used in the MRI version (i.e., bicycle, car, fish, hand, horse, house, sailboat, scissors, telephone, train, and soccer ball). The child is required to press the space bar every time he or she sees a picture that is *not* a soccer ball. They are required to withold responding everytime a soccerball is presented. The duration of a single K-CPT run is 7.5 minutes. Each administration of the K-CPT comprises 5 blocks, with each block consisting of a 20-trial sub-block featuring 1500ms inter-stimulus intervals (ISIs) followed by another 20-trial sub-block with 3000ms ISIs, resulting in a total of 200 trials. The designated time for presenting stimuli is 500 milliseconds. The K-CPT was administered by trained examiners using a standardized script and protocol. The K-CPT 2nd edition was validated on a sample of four- to seven-year-old children including children diagnosed with ADHD. The scoring software generates *T*-scores and percentiles for several variables, including commission errors, omission errors, hit RT, and hit RT variability.

## 4| DATA ANALYSIS

### 4.1| T1-weighted post-processing

T1-weighted images were visually inspected for quality control and rated on a seven-point scale, with a score of 4 representing no movement artifact and 1 indicating substantial movement artifact. The average rating for the T1-weighted images analyzed in this study was 3.712 (*SD* = 0.518). After visual inspection, the T1-weighted MRI images underwent post-processing using FreeSurfer v7.0. Any errors detected during the segmentation of grey and white matter or the subcortical segmentation and cortical parcellation were manually edited according to recommended protocols [61], and the brains were reprocessed until they met the acceptable quality control standards. The edited brains were then used in the BOLD EPI processing stream.

### 4.2| Movement

Excessive movement was defined as a framewise displacement (FD) greater than 0.9mm between successive TRs, based on [62], with participants excluded if the percentage of TRs that were censored exceeded 15%. Included and excluded groups were statistically different in terms of ADHD symptomology *t* (287.75) = −3.52, *p* < 0.0005. This resulted in the exclusion of 51 (34%) participants from the TD group and 78 (43%) participants from the ADHD group. After applying this exclusion criteria, the two groups did not significantly differ in terms of FD movement, *t*(120.61) = −1.659, *p* = 0.100.

### 4.3| BOLD EPI post-processing

We used fMRIprep [63] to post-process the MRI data. T1-weighted structural volumes were corrected for intensity non-uniformity (N4BiasFieldCorrection) [64] and skull-stripped (antsBrainExtraction from Advanced Normalization Tools; ANTS). Functional data were motion-corrected using MCFLIRT (from FSL) [65] and slice-time corrected to the middle of each TR using 3dTshift (from AFNI) [66]. Functional images were co-registered to corresponding T1-weighted images using boundary-based registration [67] with nine degrees of freedom via FreeSurfer. The motion correction transformations, distortion correction warp, functional to anatomical transformation, and anatomical to template warp were all concatenated and applied in a single step using Lanczos interpolation (from ANTs) [68].

From this point we employed two pipelines, one in the Adolescent Brain and Cognitive Development (ABCD) Atlas space, and the other in the original participant space. In the individual space, we made use of the individualized parcellation and segmentation maps generated by FreeSurfer to examine the association between performance on the K-CPT outside the scanner and brain activation during the scanning portion of the study.

We also conducted a voxel-wise analysis in the ABCD Atlas space. This atlas is normed on a large child sample from the ABCD study [60] which avoids warping to an adult template, and allows for examination of subcortical structures important for this study. To do this we warped each brain to the atlas space using nonlinear registration in ANTs. Spatial smoothing in the volume space was applied for the voxel-wise analysis only (AFNI 3dmerge; 4mm FWHM kernel).

For both pipelines, the following steps were implemented: 1) FD was calculated for each scan, and a censor file was established implementing a 0.9mm cutoff. The first three TRs of each scan were also censored to allow the MR signal to reach a steady state; 2) prior to the general linear model, percent signal change was calculated on the native time series; 3) the general linear model was employed to model the degree of BOLD activity during the K-CPT task against a resting baseline (fixation) using AFNI 3dDeconvolve. The peak amplitude was set to 1. In addition to the stimulus timing predictor, we included in the model the censor file for movement censoring, polynomial drift predictors, six movement parameters, and signal from CSF and white matter. The output of this last step included, for each voxel, beta values representing percent signal change over resting baseline, and their associated t-statistics.

Second-level group analyses were conducted in the volume space to assess the difference in brain activity during the K-CPT across groups (ADHD vs TD). The group analysis was conducted on the level-1 beta weights using Fast and Efficient Mixed Effects Algorithm (FEMA) [69], which is optimized for the ABCD brain atlas. The design matrix was set up to examine the main effect of Diagnostic Group on voxel-wise beta estimates of activation, and included age, sex assigned at birth, parental education, and movement (in number of censored TRs) as covariates. Statistical parametric maps were thresholded at a single voxel threshold of *p* < .005. Simultaneously, threshold free cluster enhancement [70] (using FEMA) was applied to the same maps. This method enhances cluster-like structures without defining clusters in a binary way. Using this method, we identified a mask of enhanced clusters on unthresholded data, and applied that mask to thresholded statistical parametric maps of each respective comparison.

## 5| REGION OF INTEREST ANALYSIS

We were primarily interested in identifying regions that are sensitive to the K-CPT task. Based on prior literature, we expected the anterior insula, inferior frontal gyrus (*pars opercularis, triangularis, orbitalis*), superior parietal lobule, anterior cingulate, and superior frontal gyrus medial wall (SMA and pre-SMA) to be the regions most likely to be involved in this task. We identified these regions (bilaterally) anatomically on individual cortical surfaces based on manual refinement of the automatic FreeSurfer parcellation [71, 72].

To assess the association between K-CPT performance measured outside the scanner, and activity in our defined ROIs, multiple regression analyses were conducted using R (v. 4.1.3;https://cran.r-project.org/). Activity for each ROI was extracted by averaging the hemodynamic response estimates (betas) for each participant for each defined ROI in the volume space (FreeSurfer aparc+aseg using the Destrieux atlas [73], based on the anatomical conventions of Duvernoy [74]). Data were missing for all four K-CPT variables measured outside the scanner (27% were missing for commissions, ommissions, and Hit RT; 35% were missing for RT Variability). In order to deal with missing data for these outcome measures, we used multiple imputation with Multivariate Imputation by Chained Equations (MICE) in R (package `mice`). Twenty imputation sets were defined, and the data were pooled during model estimation according to Rubin’s rules [75].

In addition to dealing with missing data, we also downweighted outlying values using robust regression. Outliers were thus downweighted using a Huber loss function in the regression model (R function `rlm`; [76, 77, 78]. In addition to standardized *β* and associated statistics, semipartial *r* effect size (*r_sp_*) of the predictors of interest are also reported. This measure is the correlation between the predictor and the outcome unique from the other variables in the model, and its size can be understood in the same way as the Pearson correlation.

## 6| RESULTS

### 6.1| Behavioral Compliance and Responses to the Task Inside the Magnet

We examined behavioral responses recorded by the button box for children completing the task during scanning. While we found group differences in RT standard deviation (*t* (93) = −2.084, *p* = 0.039, *β* = −0.21, *B* = −42.38, 95% Confidence Interval *B* = −82.77 to −1.99), we found no differences in terms of commissions, omissions, or mean RT (all *p* > .05). This suggests children in both groups actively engaged in the task, which is also supported by the baseline activation maps reviewed below.

### 6.2| Whole Brain Analysis

Figure 2 demonstrates that the K-CPT, as implemented, effectively engages the expected inhibitory control network, as discussed in the Introduction. The data, projected onto the average cortical surface representation of the sample, reveals significant activity above the resting baseline in various brain regions. These include bilateral visual and left motor cortex, bilateral anterior insula linked to visual attention, right middle/superior frontal cortex and parietal regions associated with working memory, midline cingulate, pre-SMA, and SMA regions involved in motor planning and execution, as well as subcortical thalamic and striatal regions related to motor execution. Remarkably, these network activations are consistently robust across groups, indicating that the participants engaged with the task as expected. They are also consistent with meta-analytic results of Go/NoGo tasks [79], as seen in Figure 2 (bottom).

**FIGURE 2.**
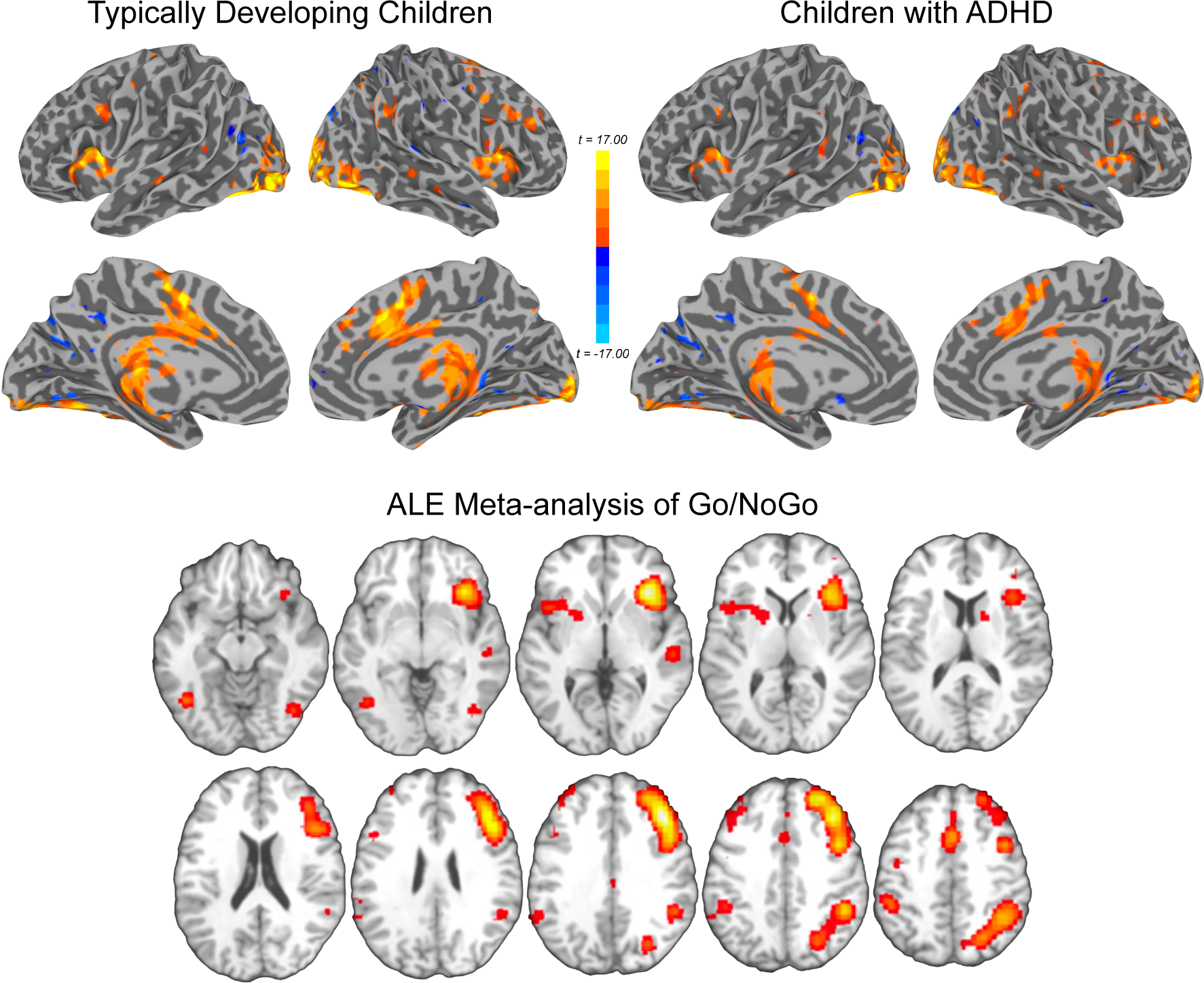
Cortical activation maps for inhibitory control. Top: Results of the whole brain analysis for *K-CPT > Resting Baseline* for both Typically Developing Children (left) and Children with ADHD (right). Results are projected to the cortical surface representation of the average sample brain (*p* < .005, cluster corrected). Bottom: For comparison, results from a meta-analysis of Go/NoGo tasks showing comparable activation. Modified from Figure 1 of Swick, D., Ashley, V., and Turken, A. U., (2011). Are the neural correlates of stopping and not going identical? Quantitative meta-analysis of two response inhibition tasks. *NeuroImage, 56*, 1655-1665.

With the robust activation of the anticipated networks established across groups, our focus shifted to investigating group differences in these networks. Figure 3 illustrates the outcomes of the TD vs. ADHD group comparison. Notably, the TD group exhibited greater activation in regions of the inhibitory control network, including the left inferior frontal gyrus and anterior insula, right pre-SMA and anterior cingulate cortex, and bilateral caudate. Intriguingly, one cluster was found in the lateral orbitofrontal cortex, which is not typically associated with inhibitory control. On the other hand, the ADHD group demonstrated greater activity than the TD group, although mostly outside the putative inhibitory control network, involving regions such as bilateral cuneus, right postcentral, and supramarginal gyrus.

**FIGURE 3.**
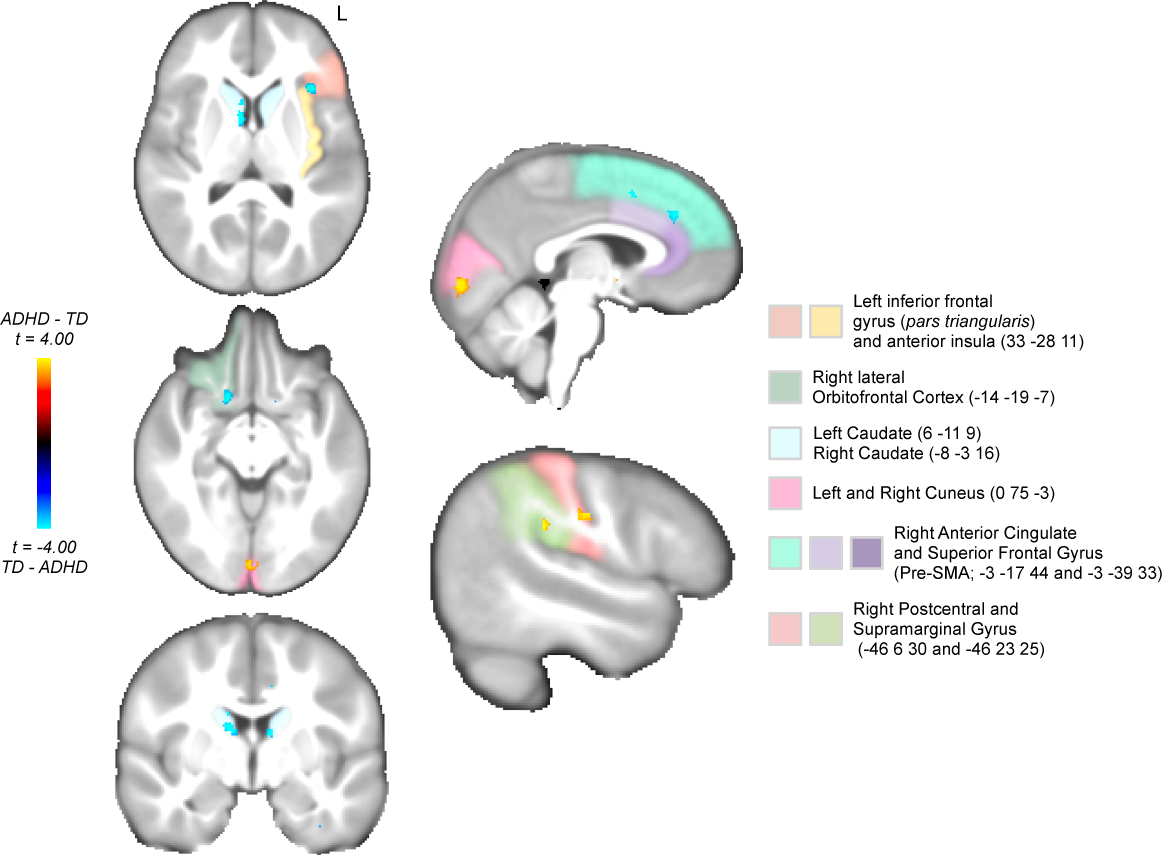
Differences in activity between ADHD and TD groups are overlayed on the ABCD group atlas. Activity favoring the TD group (shown in dark-blue-to-light-blue spectrum) was found in some regions of the putative inhibitory control network, including left inferior frontal gyrus and anterior insula, right pre-SMA and anterior cingulate, and bilateral caudate nucleus. One cluster was also found in lateral orbitofrontal cortex, not typically associated with inhibitory control. Activity favoring the ADHD group was also found in regions outside the putative inhibitory control network (shown in red-to-yellow spectrum, in bilateral cuneus, and right postcentral and supramarginal gyrus). All comparisons are *p* < .005, cluster corrected. Coordinates are reported in ABCD atlas space.

### 6.3| ROI Analysis

Next, we examined the association between brain activity in our identified ROIs and behavioral measures of the computerized K-CPT administered outside the MRI scanner. The model included diagnostic group and the group by brain interaction, in addition to sex assigned at birth, age, parent education, and movement in the scanner as covariates of non-interest.

Before movement exclusion, group differences (controlling for age, sex assigned at birth, parent education, and movement) were found for commissions (*t* (277) = −3.31, *p* = 0.001, *β* = −0.19, *B* = −3.70, 95% Confidence Interval *B* = −5.90 to −1.50), omissions (*t* (277) = −6.37, *p* < 0.0001, *β* = −0.35, *B* = −11.51, 95% Confidence Interval *B* = −15.07 to −7.95), and RT Variability (*t* (277) = −3.68, *p* < 0.0005, *β* = −0.23, *B* = −6.10, 95% Confidence Interval *B* = −9.37 to −2.83), but not for Hit RT (*t* (277) = −1.75, *p* = 0.08, *β* = −0.10, *B* = −2.05, 95% Confidence Interval *B* = −4.35 to 0.24). After movement exclusion, group differences were not found for commissions (*t* (75.36) = 1.42, *p* = 0.16, *β* = 0.30, *B* = 3.00, 95% Confidence Interval *B* = −1.20 to 7.21) or Hit RT (*t* (72.86) = 1.10, *p* = 0.28, *β* = 0.25, *B* = 2.31, 95% Confidence Interval *B* = −1.88 to 6.51), but were found for omissions (*t* (65.03) = 2.54, *p* = 0.013, *β* = 0.55, *B* = 8.55, 95% Confidence Interval *B* = 1.84 to 15.27) and RT Variability, (*t* (50.21) = 2.64, *p* = 0.011, *β* = 0.61, *B* = 8.11, 95% Confidence Interval *B* = 1.95 to 14.28).

Only one brain region was associated with K-CPT performance outside the magnet. That is, activation in the right short gyrus of the insula was associated with both Hit RT (*t* (100.27) = −2.05, *p* = 0.04, *β* = −0.28, *B* = −16.70, 95% Confidence Interval *B* = −32.85 to −0.55) and RT Variability (*t* (96.56) = 3.90, *p* = 0.014, *β* = −0.33, *B* = −26.96, 95% Confidence Interval *B* = −6.21 to −0.05; see Figure 4). Although there was a trend (*p* = .059), there was no significant Diagnostic Group by Brain interaction in either region, and neither main effect survived a FDR correction for multiple comparisons across brain regions, but we report the effect here to be comprehensive. Our interpretation is appropriately tempered in the Discussion.

**FIGURE 4.**
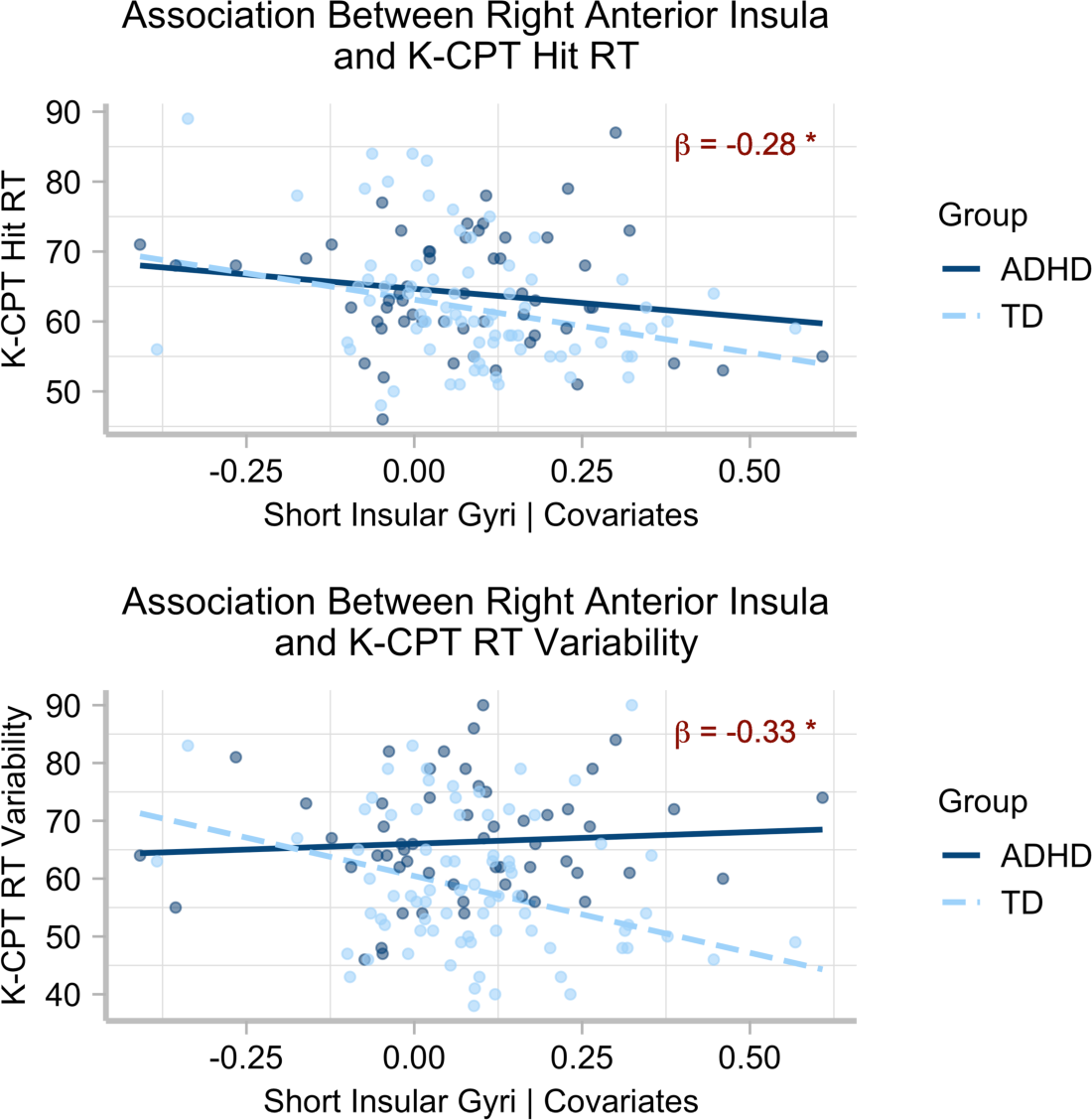
Association between right anterior insula activity and Kiddie Continuous Performance Test (K-CPT) scores, controlling for age, sex assigned at birth, family education, and movement in the scanner. Main effects were found for Hit RT and RT Variability measures. No significant Diagnostic Group by Brain interaction effects were found, but Diagnostic trends are shown for reference. *β* is the standardized regression slope for the main effect. * *p* < .05, uncorrected for multiple comparisons.

## 7| DISCUSSION

Behavioral continuous performance paradigms have demonstrated good sensitivity in distinguishing very young children with ADHD from typically developing (TD) children [37]. However, their use in understanding the neural circuitry underlying inhibitory control in these children is rare. In fact, fMRI paradigms investigating inhibitory control networks in very young children are scarce overall, despite their significance in mapping functional activation differences between ADHD and TD children. Developing such paradigms presents an opportunity to elucidate the neurobiology of inhibitory control impairments observed in children with ADHD and establish a framework for future investigations into the development of these networks in both TD children and those with ADHD. To address this objective, we studied a cohort of 56 children diagnosed with ADHD and 78 TD children, all aged 4- to 7-years. We used a modified version of the K-CPT during BOLD fMRI and assessed their performance on the clinically established and standardized K-CPT outside the MRI scanner. Our findings revealed the following: 1) the modified K-CPT effectively elicited robust activity in established inhibitory control networks for both groups; 2) group comparisons yielded small clusters of activity differences, primarily in brain regions known to be involved in inhibitory control (e.g., inferior frontal gyrus, anterior insula, dorsal striatum, medial pre-SMA, and cingulate cortex); and 3) increased activity in the right anterior insula was associated with reduced Hit RT and RT Variability on the K-CPT administered outside the MRI environment. These results highlight three key points: (1) the modified K-CPT was effectively adapted for use in the fMRI setting with very young children, successfully eliciting a robust response in expected regions for both groups; (2) the paradigm exhibited modest sensitivity to group differences in inhibitory control network regions; and (3) the paradigm captured activity associated with performance on the standard K-CPT administered outside the fMRI environment, although this effect did not survive statistical correction for multiple comparisons across the number of ROIs. Further detailed discussion of these specific results follows below.

### 7.1| The modified K-CPT elicits a robust response in regions of the inhibitory control network for both groups

The modified K-CPT elicited a robust response in brain regions of the inhibitory control network for both groups. This is consistent with a number of fMRI CPT and inhibitory control studies in adults [80, 81, 82, 83, 84]. For example, in adults these task paradigms elicit activity in brain regions associated with initiating the response (i.e., “Go” process), such as motor and premotor cortex, dorsal striatum, pallidum, and thalamus. The “Stop” process recruits right inferior frontal cortex, pre-SMA, anterior cingulate, anterior insula, and parts of the basal ganglia, including the subthalamic nucleus. In Aron and colleagues’ model of inhibitory control, right inferior frontal cortex and pre-SMA collaborate with basal ganglia circuits to initiate and implement action plans, with the subthalamic nucleus as the terminal target via hyperdirect pathways from both inferior frontal cortex and pre-SMA [85]. As expected, the baseline response was robust for both groups in these regions. It is particularly striking to note that the activation profiles were largely symmetric bilaterally, with especially robust responses in bilateral anterior insula, middle frontal gyrus, and SMA, pre-SMA, and anterior cingulate cortices. Investigation of the ALE maps from published research (Figure 2 bottom) shows that this is expected for these regions, based on studies in adults. Establishing that this paradigm elicits robust above-baseline activity is critically important for examining group differences. We can now turn to these group differences.

### 7.2| Direct group comparisons yielded activity differences in brain regions putatively involved in inhibitory control, but brain activation associations with performance outside the scanner did not differ across groups

We turn first to the elicited activity in anterior insula, which often extends into the neighboring inferior frontal gyrus (e.g., see ALE maps in Figure 2). Robust activity in the anterior insula in response to inhibitory control demands has been a focus of inquiry in recent years. It has been observed that the anterior insula is often the site of peak activation during inhibitory control paradigms [79, 86]. However, it is worth noting that the anterior insula is absent in some prominent models of inhibitory control [30, 23], although others have shown interest in defining its function more specifically. For instance, Molnar-Szakacs and Uddin [87] proposed that the anterior insula serves as a “gatekeeper of executive control,” citing its anatomical and functional associations with other regions and networks involved in executive function. This notion is supported by meta-analyses of both SST and Go-NoGo paradigms, which consistently demonstrate bilateral activation of the anterior insula, along with the pre-SMA and SMA [79]. In fact, Swick and colleagues [79] noted “results clearly demonstrate the importance of bilateral anterior insular regions and medial BA 6 (SMA/pre-SMA) for successful performance in response inhibition tasks.” Finally, a number of studies report that anterior insula is also deferentially recruited in response inhibition tasks by youth with ADHD [88, 89, 90, 91, 92], showing it’s potential sensitivity to ADHD symptomology.

Typically, inhibitory control paradigms emphasize the importance of the right hemisphere inferior frontal gyrus and anterior insula [93]. But in a direct comparison between the groups, we found that only *left* anterior insula extending into inferior frontal gyrus showed greater activation for the TD group relative to the ADHD group. We do note that at a reduced threshold (*p* < .01), the right insula/inferior frontal gyrus also showed the same activation pattern, but this did not survive the stricter threshold. In studies with adults, both left and right anterior insula are associated with inhibitory control [94] and error processing [95], with stopping efficiency in inhibitory control paradigms and general accuracy in those paradigms [96], and differences in both left and right insula are found for adults with ADHD [97]. Boehler and colleagues [96] interpreted left insula activity in inhibitory control paradigms to indicate general attention modulation, which is consistent with the role for this region ascribed by Molnar-Szakacs and Uddin [87]. It is possible the difference in activity across groups is indicative of this general attention modulation. Indeed, in a study of eighty adolescents (49 with ADHD), increasing ADHD symptom severity was associated with decreased recruitment of left anterior insula during a Go/No-Go task [88]. They likewise interpreted this association as reflecting the general attentional demands of the task, although it is difficult to tease apart the distinction between attentional load and response control in the present paradigm [98].

The right lateral frontal cortex extending into insula is also an important node in the inhibitory control network, and the regions are often coactivated in many inhibitory control paradigms, making their roles in such tasks difficult to distinguish [99]. Indeed, in the present study, although the group difference was most notable in left insula/inferior frontal gyrus, the association between activation and K-CPT performance (namely Hit RT and RT Variability) was actually found in the *right* anterior insula (defined at the ROI level). Furthermore, there was no group by brain activation interaction, suggesting that greater activation in right anterior insula was associated with both faster RT, and reduced RT variability, in *both* groups. Despite the fact that the reported association did not survive statistical correction, the effect size, especially for RT variability, was meaningful. In the latter comparison, the *β* was −0.34, which is a sizeable associations, especially considering it accounts for a number of covariates. However, the lack of significance after correction suggests we should interpret the result with caution.

In addition to the lateral cortical regions, we also observed group differences in the bilateral caudate and medial frontal cortical regions, including the anterior cingulate and pre-SMA. These regions are crucial nodes in the neural system involved in stopping behaviors, such as the ability to withhold a response in the K-CPT [100, 85, 101, 102, 103, 96, 104]. In models of inhibitory control, pre-SMA and striatum (including caudate) play a role in the downstream selection of a competing action among a space of possible action programs. The dorsal striatum is uniquely situated as the inter-face between cortex and the rest of the basal ganglia, forming part of an indirect cortico-basal ganglia-thalamic loop involved in action selection [100]. In a study of adults with ADHD, Sebastian and colleagues [105] found less activation in right caudate for people with ADHD compared to control participants, which is in line with what we report here.

Medial frontal cortex, especially SMA and pre-SMA, are projecting nodes as part of a “hyperdirect pathway” to the subthalamic nucleus (STN) [106]. These hyperdirect projections also include M1 and lateral inferior frontal cortex, and together these regions function to influence STN activity, which via globus pallidus and thalamic activation works to suppress motor output [85, 107, 108]. Pre-SMA also likely collaborates directly with lateral inferior frontal cortex as part of this loop [85, 109]. Given that we found differences in several key nodes of this putative inhibitory control network, we could cautiously interpret the findings to indicate differences in recruitment of the whole network, which may in turn indicate group differences in the efficiency or functional dynamics in response to inhibitory control demands. However, we should also be careful here to not overstep the limits of the BOLD paradigm as it relates to the measurement of network-level dynamics.

Other regions outside the putative inhibitory control network also showed group differences. For example, right supramarginal gyrus and postcentral gyrus, and occipital cortex, showed greater activity for the ADHD group. Right supramarginal gyrus and occipital cortex were actually identified as reliably active by Boehler and colleagues [96] in a comprehensive conjunction analysis of the stop-signal paradigm, along with regions of the putative inhibitory control network. Activation in these regions was interpreted to indicate sensory processing of the stop stimulus (occipital cortex) and attentional modulation of sensory stimuli, or bottom-up attentional recruitment (supramarginal gyrus). It is possible that less efficient processing in frontal regions of the inhibitory control network elicited greater demand on these regions involved in attentional modulation, leading to recruitment to a greater degree in the ADHD group.

### 7.3| Limitations

Although this study suggests that the modified K-CPT paradigm can be used in young children and is sensitive to group differences in regions of the inhibitory control network, as well as the association between activation in the right insula and task behavior outside the scanner, there are several significant limitations. First, movement was a substantial issue for a large portion of the sample, which was expected given the young age of the participants [110]. To address this, we implemented a strict movement cutoff to establish a reliable inhibitory control network against the resting baseline. While this cutoff aligns with optimal analytic protocols for dealing with movement [62], it is more stringent than what is sometimes used in pediatric fMRI studies, which can set cutoffs as large as a voxel or more. Consequently, we excluded a considerable number of both TD and ADHD children (see, e.g., Gaffrey et al., 2020 for comparable attrition in this age range), potentially introducing a selection bias favoring children with less severe ADHD symptoms within the ADHD group. Thus, the observed group differences may not be as pronounced as they would have been with a less strict movement criteria. In addition, this limits the external validity of the study, as the sample represents children who are able to stay still in a MRI scanner. However, employing a more lenient movement criteria would likely have compromised the reliability of the data. It is worth noting that movement poses a significant challenge in pediatric fMRI [111], as it strongly influences the initial estimation of the BOLD response due to movement-related noise [112]. While including movement as a regressor in the second-level analysis is important, it does not completely address the issue of movement in the first-level estimation of the BOLD response. One could argue that our arbitrary inclusion cutoff of 15% of the time series is overly conservative, but unfortunately, to our knowledge no established guidelines exist in the literature [113]. Therefore, we opted for caution. Consequently, the reported differences should be interpreted in light of the final analyzed sample. This sample successfully facilitated a meaningful comparison between ADHD and TD children. Notably, all ADHD children met strict diagnostic criteria as determined by dual clinicians, and we observed behavioral differences in K-CPT performance, particularly in terms of reaction time (RT) variability, which is the most sensitive K-CPT measure associated with ADHD [37]. However, group differences might have been more pronounced if children with more significant ADHD symptomatology had been included in the final sample.

A second limitation of this study as it relates to identified network differences across groups is the young age of the sample. Most investigations of children with ADHD involve older children (typically over 9-years of age) and young adults. Studies of typical children suggest that functional networks change, sometimes markedly, over development [114, 18]. For example, in a study of verbal fluency from 7-18-years, Holland and colleagues [114] reported increased left laterality of activation as a function of age. In other words, the functional network of this standard executive control task shifted substantially as children entered adolescence. Thus, it is possible that the reported differences apply generally to younger children, but may not in fact reflect activation profiles of older children with ADHD. For example, in a small sample of 6-7-year-old children during a Go/No-Go paradigm, left anterior insula was also identified as more active in control children compared to children born preterm [115]. Whether these regions remain the key nodes in older children would be most appropriately addressed in a longitudinal investigation.

### 7.4| Conclusions

To briefly summarize the findings, our research uncovered the following: 1) the modified K-CPT successfully triggered strong and expected brain activity related to inhibitory control networks in both groups of participants; 2) when comparing the two groups, we observed modest differences in brain activity primarily in regions associated with inhibitory control, including the inferior frontal gyrus, anterior insula, dorsal striatum, medial pre-SMA, and cingulate cortex; 3) heightened activity in the right anterior insula was linked to quicker and more consistent response times on the K-CPT conducted outside the MRI scanner. The findings broadly support the usefulness of this paradigm for very young children. However, they also reveal the limitations in studying children with ADHD in this age range within the MRI environment, even with careful procedures to minimize movement. This led to the need for a large initial sample size to ensure an appropriate investigative sample. Our exclusion criteria, especially for movement, might have resulted in the exclusion of children with more severe ADHD, and could limit the differences between the two groups. Therefore, we should interpret our reported results in the context of these limitations and the young age of the sample.

## Abbreviations

ADHD: Attention-Deficit/Hyperactivity Disorder

## Acknowledgements

We thank the families and children who participated, and continue to participate, in the AHEAD study, as well as staff involved in data collection.

## Author contributions statement

A.S.D and P.G. conceptualized and designed the study. A.S.D. and M.B. analyzed the data and wrote the draft manuscript. M.H., M.C., D.G. contributed to data collection and data curation. All authors reviewed and commented on the draft manuscript and all authors reviewed and approved the manuscript.

## Declaration of generative AI and AI-assisted technologies in the writing process

During the preparation of this work the authors used ChatGPT in order to improve grammar and flow of content originally-drafted by the authors. After using this tool/service, the authors reviewed and edited the content as needed and take full responsibility for the content of the publication.

